# The *Klebsiella pneumoniae* citrate synthase gene, *gltA*, influences site specific fitness during infection

**DOI:** 10.1101/568493

**Authors:** Yuang Sun, Jay Vornhagen, Paul Breen, Valerie Forsyth, Lili Zhao, Harry L.T. Mobley, Michael A. Bachman

## Abstract

*Klebsiella pneumoniae* (Kp), one of the most common causes of healthcare-associated infections, increases patient morbidity, mortality and hospitalization costs. Kp must acquire nutrients from the host for successful infection. However, the host is able to prevent bacterial nutrient acquisition through multiple systems, including the innate immune protein lipocalin 2 (Lcn2) that prevents Kp iron acquisition by sequestering the siderophore enterobactin. To identify novel Kp factors that mediate evasion of nutritional immunity during lung infection, we undertook an InSeq study using a pool of >20,000 transposon mutants administered to *Lcn2+/+* and *Lcn2-/-* mice. Comparing mutant frequencies between mouse genotypes, this genome-wide screen identified the Kp citrate synthase GltA as potentially interacting with Lcn2, and this novel finding was independently validated. Interestingly, *in vitro* studies suggest that this interaction is not direct. Given that GltA is involved in oxidative metabolism, we screened the ability of this mutant to use a variety of carbon and nitrogen sources. The results indicated that the *gltA* mutant has a distinct amino acid auxotrophy and is unable to use a variety of carbon sources. Specifically, we show that *gltA* is necessary for growth in bronchioloalveolar lavage fluid, which is amino acid-limited, but dispensable in serum, which is amino acid rich. Deletion of *Lcn2* from the host leads to increased amino acid levels in bronchioloalveolar lavage fluid, and thus abrogates the loss of *gltA* during pneumonia in the *Lcn2-/-* background. GltA was also required for gut colonization, but dispensable in the bloodstream in a bacteremia model, demonstrating that deletion of *gltA* leads to an organ-specific fitness defect. Together, this study is the first to mechanistically describe a role for *gltA* in Kp infection and provide unique insight into how metabolic flexibility impacts bacterial fitness during infection.

**Author Summary:** The bacteria *Klebsiella pneumoniae* (Kp) is an important cause of infection in healthcare settings. These infections can be difficult to treat, as they frequently occur in chronically ill patients and the bacteria has the ability to acquire multiple antibiotic resistance markers. Kp is a common colonizer of the intestinal tract in hospitalized patients, and can progress to infections of the bloodstream, respiratory and urinary tract. However, the bacterial factors that allow Kp to replicate in these different body sites is unclear. In this study, we found that the Kp citrate synthase, GltA, enables bacterial replication in the lung and intestine by enhancing the ability of Kp to use diverse nutrients, in a mechanism known as metabolic flexibility. Kp lacking GltA require specific amino acids that are abundant in blood, but not other body sites. The work in this study provides novel insight into why Kp is a successful hospital pathogen that can colonize and infect multiple body sites.

## Introduction

Extended spectrum beta-lactamase (ESBL)-producing *Enterobacteriaceae* and carbapenem-resistant *Enterobacteriaceae* (CRE) pose a serious public health threat due to their extensive antibiotic resistance. Many ESBL and CRE infections are healthcare-associated infections (HAIs), meaning they occur in long-term healthcare facilities and hospitals. *Klebsiella pneumoniae* (Kp) is an environmentally ubiquitous member of the *Enterobacteriaceae* family that can acquire antibiotic resistance genes [1], and thus, is a leading cause of ESBL-producing *Enterobacteriaceae* infections [2] and HAIs [3]. Strikingly, mortality rates in patients infected with antibiotic resistant Kp often exceed 40% [4]. Disturbingly, reports of hypervirulent clones of Kp acquiring mobile antibiotic resistance genes are becoming more frequent [5,6], posing a significant global threat to human health. As the efficacy of antibiotics diminishes and therapeutic options for patients infected with antibiotic resistant strains of Kp become increasingly limited, a better understanding of how Kp establish productive infections is necessary for the development of novel diagnostics and interventions to combat these dangerous bacteria.

To establish a productive infection, bacterial pathogens such as Kp must acquire nutrients from the host environment. Subsequently, metabolic flexibility frequently dictates the capacity of pathogens to invade different niches [7–10]. This flexibility is defined by the ability of a bacterial pathogen to categorically or conditionally acquire and utilize different metabolites. For example, *Salmonella enterica* serotype Typhimurium uses tetrathionate as an electron acceptor, providing a fitness advantage in the gut, whereas this advantage is not conferred in the spleen due to the lack of tetrathionate [11]. Interestingly, Kp potentially exhibits diversity in metabolism and nutrient acquisition, as indicated by the ability to cause a wide range of severe infections, including pneumonia, bacteremia, urinary tract infection, and pyogenic liver abscess [12]. Additionally, infectious Kp frequently originates from sites of colonization [13–15], including the gut and nasopharynx [16,17]; however, the impact of metabolic flexibility on Kp pathogenesis has not received significant attention.

Metabolites necessary for niche invasion by pathogens can be acquired directly from the host, through the metabolic activity of other microorganisms within the host microbiome, or by *de novo* synthesis, and limitation of access to these nutrients by the host is a universal means of impeding niche invasion by bacterial pathogens [18–21]. For example, iron (Fe) is critical for niche invasion and subsequent pathogenesis. The host sequesters ferrous iron (Fe^2+^) by complexing with heme and ferric iron (Fe^3+^) by complexing with transferrin, ferritin, or lactoferrin [18]. To overcome these complexes, bacterial pathogens such as Kp encode a variety of proteins and small molecules to harvest sequestered iron, including the family of low molecular weight chelators known as siderophores [22]. Consequently, the host further prevents iron acquisition by sequestering bacterial siderophores with innate immune molecules such as Lipocalin 2 (Lcn2). Lcn2 specifically binds the bacterial siderophore enterobactin [23]; however, highly pathogenic bacterial strains encode alternative siderophores that circumvent this activity. This strategy has been observed during Kp lung infection, wherein the presence of the alternative Kp siderophore yersiniabactin is sufficient to overcome the effects of Lcn2 [24]. Interestingly, Lcn2-bound Kp enterobactin has immunomodulatory effects [25], suggesting that the impact of Lcn2 during Kp infection is not limited to sequestration of enterobactin.

Beyond the well-characterized interaction between Lcn2 and siderophores during Kp infection, little is known about other Lcn2-interacting Kp gene products in the context of nutrient acquisition. To discover Kp genes that are conditionally essential in the presence of Lcn2, we undertook an InSeq experiment comparing lung infection in *Lcn2+/+* and *Lcn2-/-* murine backgrounds. This revealed multiple conditionally essential genes, including the citrate (Si)-synthase gene *gltA. In vitro* studies indicate that the interaction between GltA and Lcn2 is indirect. Further analysis revealed that deletion of *gltA* dramatically reduces metabolic flexibility, leading to arginine, glutamine, glutamate, histidine, and proline auxotrophy and a severe limitation of carbon utilization. This limitation of metabolic flexibility is partially complemented during lung infection by deletion of *Lcn2* due to an increase in amino acid concentrations in bronchioloalveolar lavage fluid, and GltA is dispensable for growth in amino acid-rich environments, such as serum. Finally, using multiple murine models of infection, we show that GltA is not necessary for bacteremia, but is necessary for gut colonization independent of the presence of *Lcn2.* Together, these data reveal how Kp metabolic flexibility, conferred by the citrate (Si)-synthase GltA, impacts compartmentalized fitness during infection.

## Results

To comprehensively identify novel LCN2-interacting Kp gene products during Kp lung infection, we employed a previously described transposon library in the Kp strain KPPR1 [26,27]. To this end, *Lcn2+/+* and *Lcn2-/-* mice [28] were retropharyngeally inoculated with 1.4×10^6^ CFU of a pool of ~25,000 transposon mutants (Fig S1A). Twenty-four hours after inoculation, mice were euthanized, lungs were collected and homogenized, and total lung CFU were collected for DNA extraction and InSeq analysis, as previously described [26]. After filtering, each sample had greater than 50 million reads corresponding to greater than 20,000 unique transposon insertions inside of open reading frames (Dataset S1). To identify LCN2-interacting Kp genes, the number of transposon insertion reads within each gene were compared between the *Lcn2+/+* and *Lcn2-/-* lung pools (mean reads per gene were 169 and 156, respectively). The log *Lcn2+/+:Lcn2-/-* insertion ratio of 1677 genes were significantly enriched or depleted after correction for multiple comparisons, including *entB* that is required to synthesize both enterobactin and the Lcn2-evading siderophore salmochelin (Dataset S1). A total of 49 genes had a fitness index greater or less than 3 standard deviations from the mean log *Lcn2+/+:Lcn2-/-* insertion ratio (Fig S1B, Dataset S1) and 43 of these genes were considered significant after correction for multiple comparisons (Fig S1C, Dataset S1, in bold). Of the 43 interrupted genes, 16 were enriched in the *Lcn2+/+* lung pool, indicating enhanced fitness when *Lcn2* is present, and 27 were enriched in the *Lcn2-/-* lung pool, indicating enhanced fitness when *Lcn2* is absent. Interestingly, the most common molecular function of genes enriched in both backgrounds was metabolism [29], accounting for 10 of 43 genes (23.3%), followed by membrane transport (6 of 43, 14.0%), including the putative siderophore transport system ATP-binding protein YusV, transcription factors (4 of 43, 9.3%), two-component systems (3 of 43, 6.9%), DNA repair and recombination (2 of 43, 4.7%), protein export (1 of 43, 2.3%), and quorum sensing (1 of 43, 2.3%). The molecular function of 16 (37.2%) of these genes has not been characterized. Three gene *(gltA, envZ*, and VK055_4417) enriched in the *Lcn2-/-* background were previously identified as fitness factors during Kp lung infection [26] (Dataset S1, in italics). Five genes displayed a greater than 2-log enrichment in the *Lcn2-/-* lung pool, and one, VK055_1802, had a *P* value less than 10^−300^ (Fig S1C-D, Dataset S1). VK055_1802 is annotated as *gltA*, which encodes the citric acid cycle enzyme citrate (Si)-synthase [30]. Together, these data indicate that metabolism is critical for the interaction between KPPR1 and the host during lung infection.

To confirm the role of *gltA* during lung infection and its interaction with Lcn2, we constructed an isogenic *gltA* mutant in the KPPR1 background using the Lambda Red recombinase system [31] and complemented the gene in trans. The KPPR1Δ*gltA* strain displayed no growth defect compared to WT KPPR1 in nutrient-rich media (Fig S1E). This mutant was mixed 1:1 with its WT parent strain, then inoculated retropharyngeally in both *Lcn2+/+* and *Lcn2-/-* mice. As observed with InSeq data, the KPPR1Δ*gltA* displayed a significant fitness defect compared to the WT KPPR1 strain in the *Lcn2+/+* lung that was partially alleviated in the *Lcn2-/-* lung (Fig 1A). Given that citrate can act as a weak siderophore [32] and is a component of more complex siderophores [33,34], we hypothesized that deletion of *gltA* may inhibit novel stealth siderophore activity in the presence of Lcn2. To test this hypothesis, we grew a variety of KPPR1-derived strains in RPMI 10% resting human serum ± recombinant human Lcn2. The WT KPPR1 strain was not affected by the presence of Lcn2; however, the siderophore-null KPPR1Δ*entBΔybtS* strain [35] was unable to grow in Lcn2-free conditions, indicating the importance of siderophore function for KPPR1 growth. The enterobactin-dependent KPPR1Δ*iroAΔybtS* strain [36] was able to grow in Lcn2-free conditions, but unable to grow in the presence of Lcn2, validating the antagonistic relationship between enterobactin and Lcn2 (Fig 1B). Surprisingly, deletion of *gltA* had no impact growth in the presence of Lcn2 (Fig 1B), suggesting that the relationship between *gltA* and Lcn2 during lung infection is indirect.

**Fig. 1.**
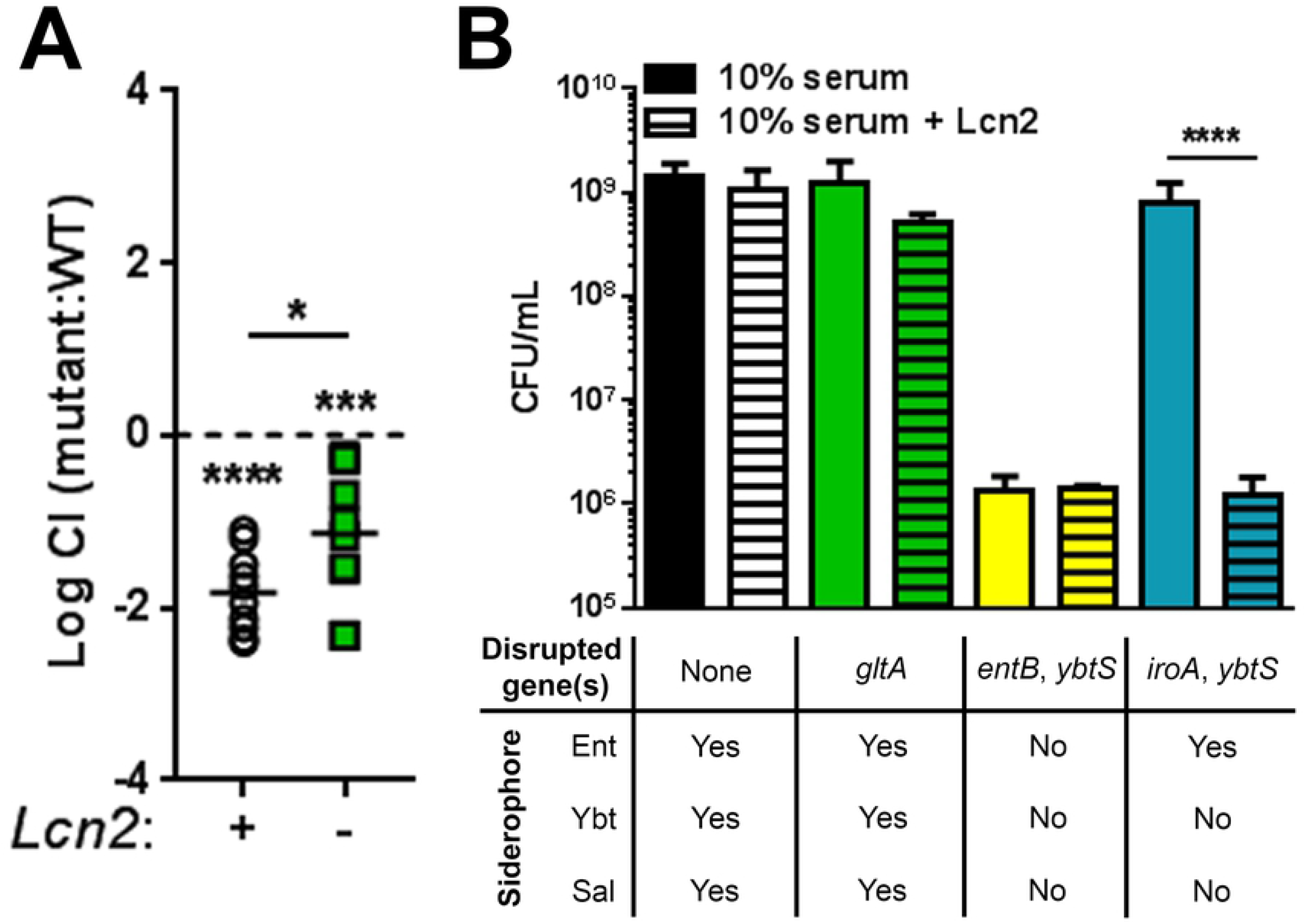
The Kp citrate synthase, *gltA*, interacts indirectly with Lcn2 during lung infection. (A) A Lambda Red recombinase mutant of *gltA* (VK055_1802) was constructed and used to validate InSeq findings. C57BL/6J mice or isogenic *Lcn2-/-* mice were retropharyngeally inoculated with approximately 1×10^6^ CFU of a 1:1 mix of WT KPPR1 and KPPR1ΔgltA. Lung bacterial burden was measured after 24 hours, and log competitive index of the mutant strain compared to the WT strain was calculated for each mouse strain (n = 10, mean displayed, \**P* < 0.05, \*\*\**P* < 0.0005, \*\*\*\**P* < 0.00005, one-sample *t* test or Student’s *t* test). (B) WT KPPR1 and various isogenic mutants were grown in RPMI + 10% (v/v) heat-inactivated resting human serum ± purified recombinant human Lcn2 overnight, then total CFU was enumerated by dilution plating on selective media (n = 3-4, mean displayed ± SEM, \*\*\*\**P* < 0.00005, Student’s *t* test).

Given that citrate (Si)-synthase performs an irreversible oxidation in the citric acid cycle, we next postulated that the relationship between *gltA* and Lcn2 is due to disruption of the TCA cycle. In addition to oxidative metabolism, the TCA cycle provides a number of key carbon skeletons for the biosynthesis of amino acids. To determine if loss of GltA affects Kp biosynthetic capabilities, we performed an unbiased screen of carbon and nitrogen sources using the BioLog system to identify conditions differentially permissive to KPPR1ΔgltA growth [37]. Both the WT KPPR1 and KPPR1ΔgltA were cultured under 288 different conditions in triplicate, and growth was measured after 24 hours (Fig 2A, Dataset S2). These experiments revealed 130 conditions in which WT KPPR1 significantly outgrew KPPR1ΔgltA, two conditions in which KPPR1ΔgltA significantly outgrew WT KPPR1, nine conditions where both strains grew equally well, and 147 conditions that did not support growth (Fig 2A, Dataset S2). These data revealed that deletion of *gltA* in KPPR1 resulted in arginine, glutamine, glutamate, histidine, and proline auxotrophy as indicated by the ability of these amino acids to support growth of the KPPR1ΔgltA strain (Fig 2B, Dataset S2). Additionally, some dipeptides containing these residues were partially or fully able to support growth of the KPPR1ΔgltA strain, whereas dipeptides without these residues do not (Fig 2C, Dataset S2). To confirm these findings, we replicated these experiments by growing the WT KPPR1, KPPR1ΔgltA, and pGltA complemented strains in minimal medium (M9) containing glucose. When glucose is the sole carbon source, the KPPR1ΔgltA strain is unable to grow but the addition of 10 mM glutamate fully restores growth (Fig 3A) and expression of *gltA* from a plasmid complemented the mutant. This auxotrophy was fully or partially complemented by addition of 10 mM glutamine, 2-oxoglutaric acid, and proline (Fig S2). Finally, the restoration of KPPR1ΔgltA growth in M9 medium containing glucose by addition of glutamate is dose-dependent (Fig 3B). These data show that deletion of *gltA* induces specific amino acid auxotrophy.

**Fig 2.**
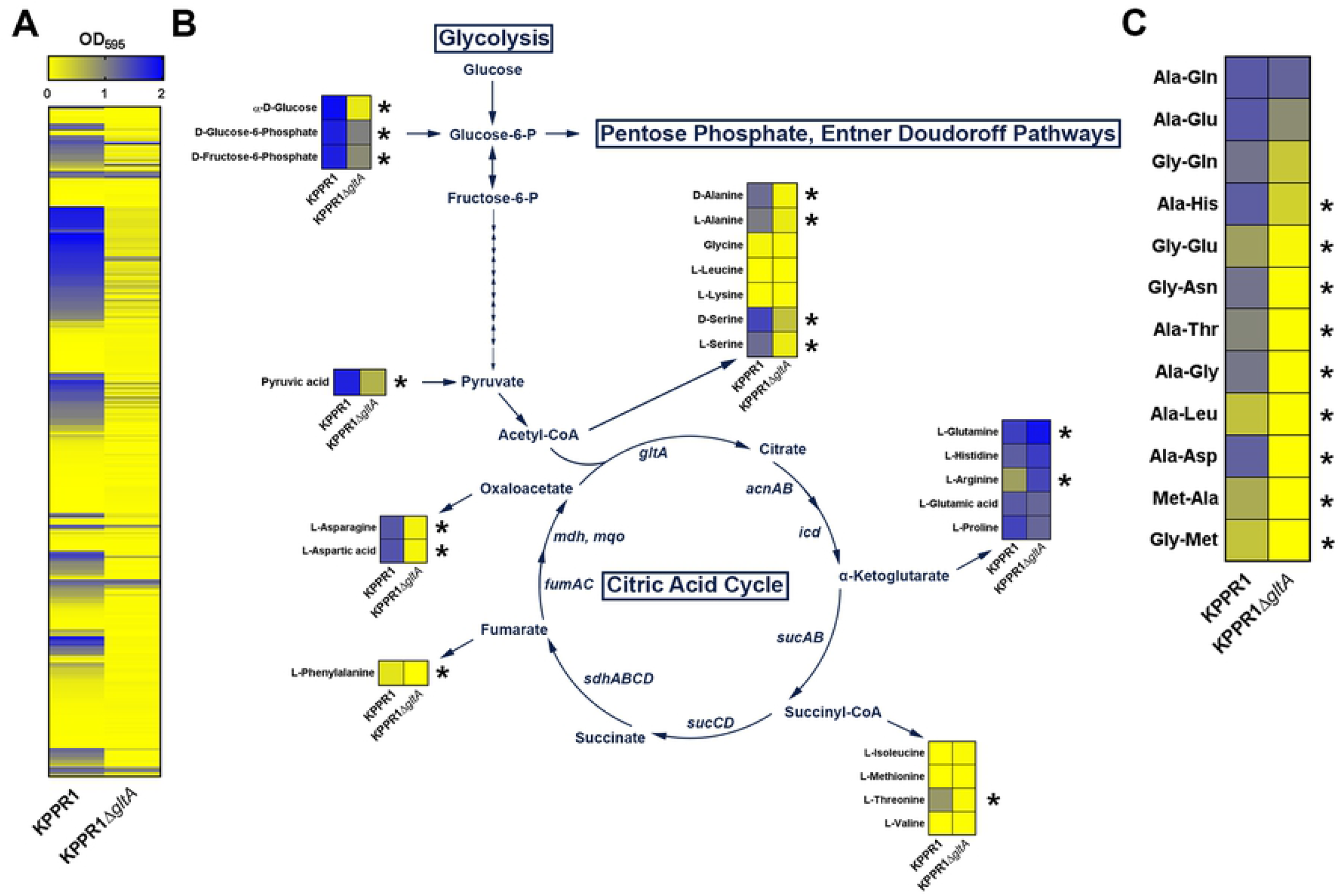
Deletion of *gltA* leads to diminished metabolic flexibility and distinct amino acid auxotrophy. (A) Heatmap summarizing BioLog Phenotype Microarray analysis of WT KPPR1 and KPPR1ΔgltA growth in in 288 carbon and nitrogen limited growth conditions indicates multiple conditions that sustained growth of WT KPPR1 but not KPPR1ΔgltA. (B) A subset of growth conditions summarizing glycolysis and non-essential amino acid biosynthesis that indicates a distinct amino acid auxotrophy is induced by deletion of *gltA.* Arrows from citric acid cycle intermediates indicate amino acids that utilize these intermediates for biosynthesis. (C) A subset of growth conditions summarizing dipeptide utilization that further indicates induction of a distinct amino acid auxotrophy by deletion of *gltA* (n = 3, mean displayed, \**P* < 0.05, Student’s *t* test).

**Fig 3.**
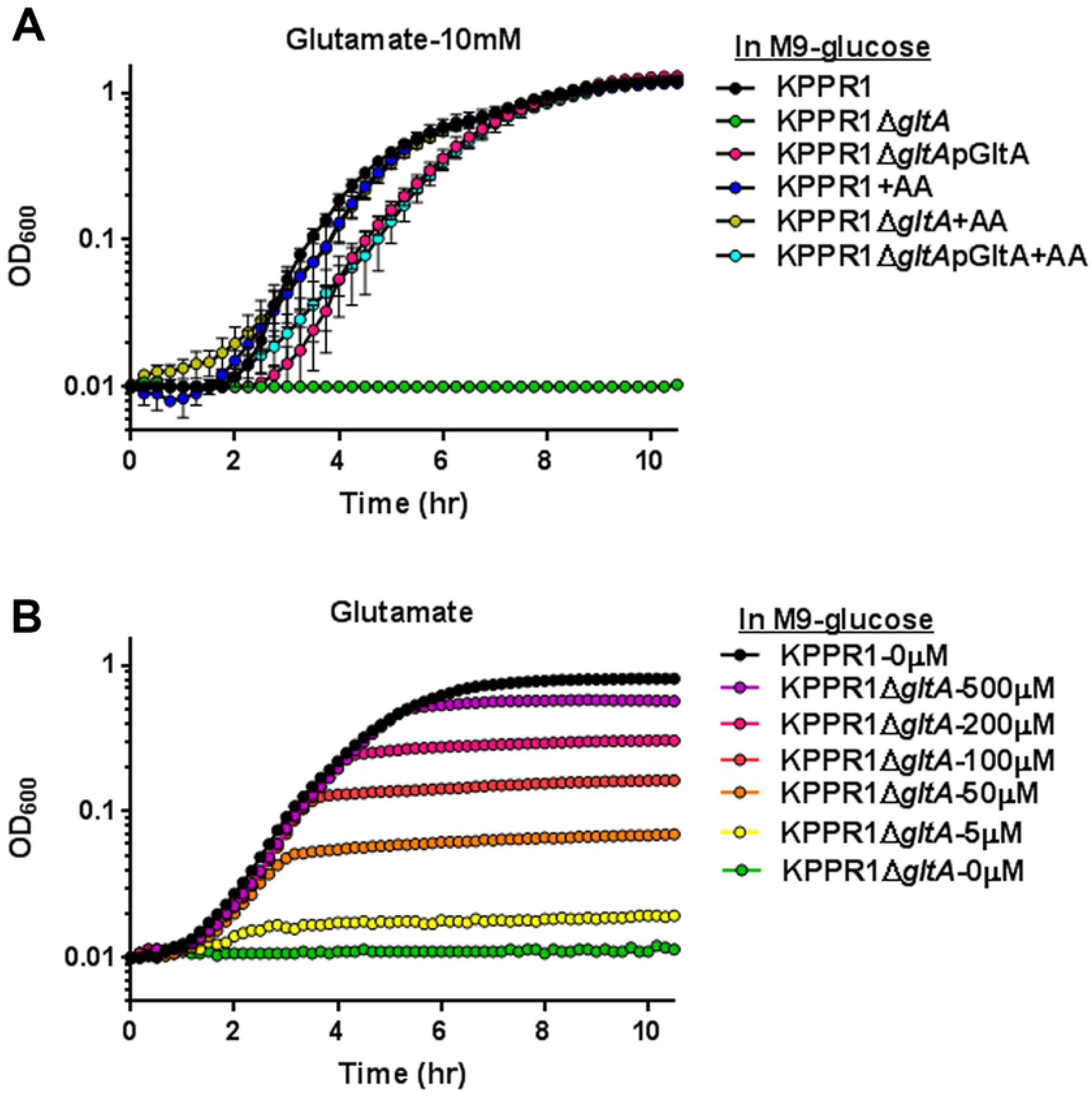
Auxotrophy due to deletion of *gltA* is functionally complimented by glutamate. (A) WT KPPR1, KPPR1ΔgltA, and KPPR1Δ*gltA* pGltA were grown in M9 minimal media + 20% glucose with 10 mM glutamate (n = 3, mean displayed ± SEM) or (B) increasing concentrations of glutamate (n = 3, mean displayed ± SEM). “+AA” label indicates addition of amino acid to growth media at concentrations indicated in graph title.

Our data indicated that the deficiency of KPPR1ΔgltA in the *Lcn2+/+* lung is related to access to specific amino acids necessary for complete growth. Bronchioloalveolar lavage fluid (BALF) is an ample nutritional source for bacteria living in the upper respiratory tract, and measurable levels the amino acids arginine, glutamate, 2-oxoglutaric acid, and proline are present in both human and mouse BALF [38–41]. To determine if the differences in KPPR1Δ*gltA* fitness in *Lcn2+/+* and *Lcn2-/-* lungs is attributable to this source of nutrient, whole BALF was collected from uninfected mice and used as a bacterial growth medium. BALF from both mouse strains sustained growth of WT KPPR1 and KPPR1Δ*gltA* (Fig 4A); however, KPPR1Δ*gltA* grew significantly less well as measured by area under curve (AUC) analysis in *Lcn2+/+* BALF compared to *Lcn2-/-* BALF and WT KPPR1 cultured in *Lcn2-/-* BALF (Fig 4B). To determine if there are inherent differences in amino acid levels in the lungs of *Lcn2+/+* and *Lcn2-/-* mice that explain these differences in growth, we used gas chromatography–mass spectrometry to determine amino acid content in BALF from uninfected mice. *Lcn2-/-* BALF contains significantly higher levels of multiple amino acids and total protein content (Fig 4C, Dataset S3). Indeed, BALF from *Lcn2+/+* and *Lcn2-/-* are fully distinguishable based on their amino acid composition (Fig 4D). This increase in amino acid and protein content likely explains the difference between KPPR1Δ*gltA* fitness in the *Lcn2+/+* and *Lcn2-/-* backgrounds by functionally complementing the loss of *gltA.*

**Fig 4.**
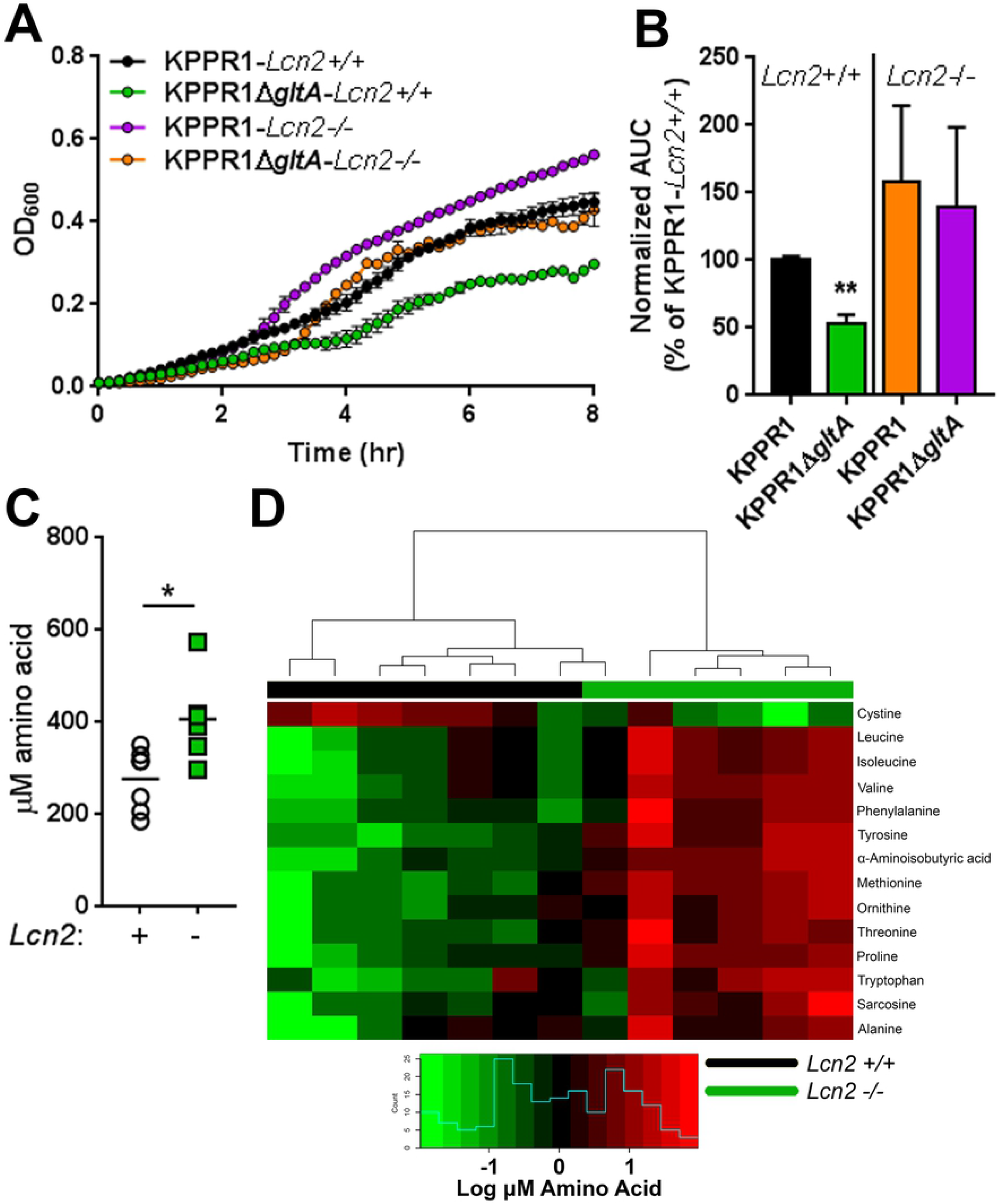
Bronchoalveolar lavage fluid from *Lcn2-/-* mice can sustain growth of KPPR1Δ*gltA* due to increased amino acid levels. (A) Murine bronchoalveolar lavage fluid (BALF) was obtained from uninfected C57BL/6J mice or isogenic *Lcn2-/-* mice, and WT KPPR1 and KPPR1ΔgltA were grown 100% BALF (n = 3, mean displayed ± SEM). (B) Area under curve (AUC) analysis was used to compare growth of WT KPPR1 and KPPR1ΔgltA in *Lcn2+/+* and *Lcn2-/-* BALF. Data are normalized to growth of WT KPPR1 in *Lcn2+/+* BALF (n = 3, mean displayed ± SEM, *\*\*P* < 0.005, ANOVA followed by Sidak’s multiple comparisons post-hoc test). (C) Total amino acid content from uninfected C57BL/6J mice or isogenic *Lcn2-/-* mice subjected to metabolomic analysis (n = 6-7, mean displayed, \**P* < 0.05, Student’s *t* test). (D) Heatmap of amino acid concentrations in BALF obtained from uninfected C57BL/6J mice or isogenic *Lcn2-/-* mice subjected to metabolomic analysis (significantly different [FDR *P* < 0.05] amino acid concentrations displayed, n > 6 mice/group). Blue histogram in inset indicates composite amino acid concentration values in heatmap matrix.

We next hypothesized that the loss of *gltA* would not affect bacterial growth in a physiologically relevant amino acid-rich environment, such as serum. To test this hypothesis, we grew the WT KPPR1 and KPPR1Δ*gltA* strains in minimal medium containing 20% heat-inactivated human serum. No differences in growth between the strains was observed (Fig 5A), and this result was recapitulated in both heat-inactivated and non-heat-inactivated murine sera (Fig S3). To determine if amino acid levels in human serum are sufficient to functionally complement the loss of *gltA*, we tested the growth of WT KPPR1 and KPPR1ΔgltA strains in minimal medium with plasma-level concentrations of glutamine, glutamate, and proline [42,43]. Indeed, serum-level concentrations of glutamine, glutamate, and proline were able to support growth of the KPPR1ΔgltA strain (Fig 5B). These data support the indication that amino acid auxotrophy induced by deletion of *gltA* is the basis of the loss of fitness observed in the amino acid poor environment of the *Lcn2+/+* lung.

**Fig 5.**
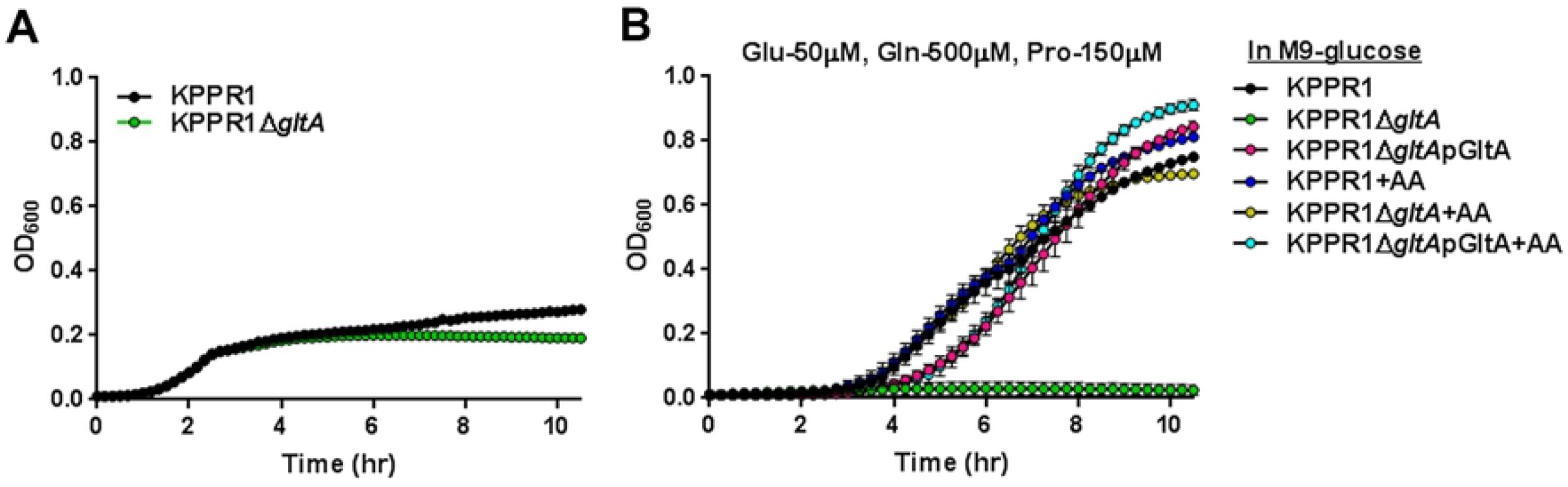
*gltA* is dispensable for growth in human serum. (A) WT KPPR1 and KPPR1ΔgltA were grown in M9 minimal media + 20% resting human serum (n = 3, mean displayed ± SEM). (B) WT KPPR1, KPPR1ΔgltA, and KPPR1 ΔgltApGltA were grown M9 minimal media + 20% glucose with physiological levels of amino acids present in human serum (n = 3, mean displayed ± SEM). “+AA” label indicates addition of amino acids to growth media at concentrations indicated in graph title.

The differential necessity of *gltA* for growth in BALF and serum led us to hypothesize that *gltA* is a fitness factor in nutritionally deplete body sites, but not in nutritionally replete body sites. To test this hypothesis, we first employed a peritoneal injection murine model of Kp infection. We injected *Lcn2+/+* and *Lcn2-/-* mice with approximately 5×10^5^ CFU of a 1:1 mix of WT KPPR1 and KPPR1ΔgltA intraperitoneally. Twenty-four hours after inoculation, mice were euthanized, and blood, liver, spleen, and lungs were collected. Solid organs were homogenized, and CFU was enumerated by dilution plating. As observed in the lung infection model, KPPR1ΔgltA was at a competitive disadvantage compared to WT KPPR1 in the *Lcn2+/+* lung, and this disadvantage was alleviated in the *Lcn2-/-* lung (Fig 6A). Consistent with *ex vivo* serum growth, KPPR1ΔgltA was not competitively disadvantaged in the blood of either mouse background but was less fit in the spleen and liver of *Lcn2+/+* mice (Fig 6A). Next, we assessed the role of *gltA* in a different nutritionally replete body site: the large intestine. The small intestine is thought to limit microbial density through absorption of nutrients; however, the large intestine is nutritionally replete, wherein bacteria utilize complex carbohydrates to sustain high population densities [44]. Results from our metabolic screen (Fig 2A, Dataset S2) indicate that the KPPR1ΔgltA strain is unable to utilize multiple sugars, thus we hypothesized that the KPPR1ΔgltA strain would be less fit in large intestine. To confirm our previous findings, we compared the growth of WT KPPR1 and KPPR1ΔgltA in a variety of conditions wherein only a single sugar was provided as a carbon source. Indeed, the KPPR1Δ*gltA* strain was unable to grow with multiple single sugars as carbon source, conditions representative of the sugars available during gut colonization (Fig S4A-K). Finally, we tested the role of *gltA* in an oral inoculation model of murine Kp infection. To this end, we gavaged *Lcn2+/+* and *Lcn2-/-* mice with approximately 5×10^6^ CFU of a 1:1 mix of WT KPPR1 and KPPR1ΔgltA. Twenty-four hours after inoculation, mice were euthanized, cecal contents were collected, and CFU was enumerated by dilution plating. In contrast to our previous experiments, we found that *gltA* is necessary for cecal colonization in both the *Lcn2+/+* and *Lcn2-/-* background (Fig 6B). Together, these data clearly display that *gltA* influences compartmentalized fitness during infection by limiting the ability of Kp to use different nutrient sources for growth during infection.

**Fig 6.**
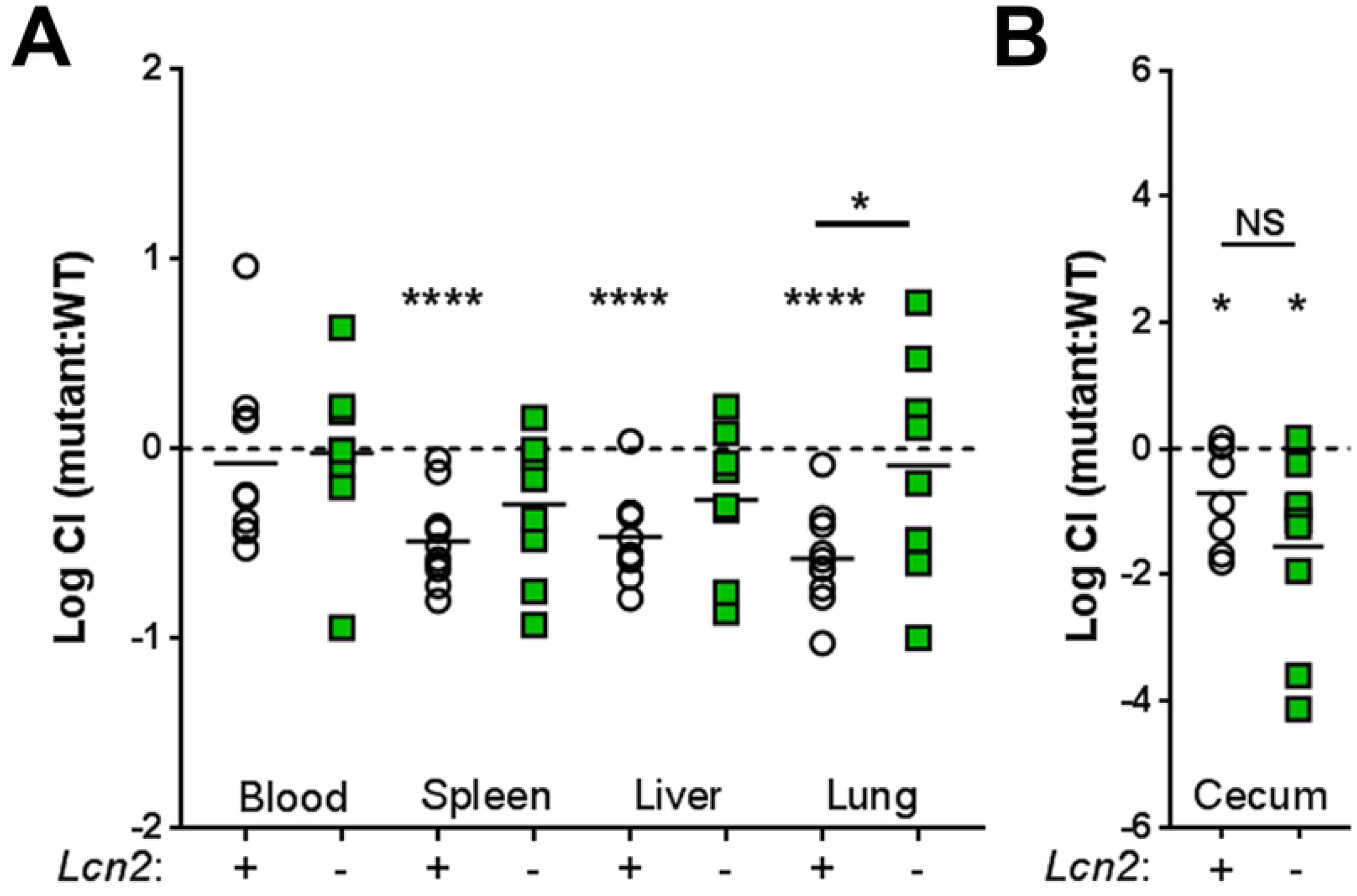
*gltA* influences compartmentalized fitness during bacteremia. (A) C57BL/6J mice or isogenic *Lcn2-/-* mice were intraperitoneally inoculated with approximately 5×10^5^ CFU of a 1:1 mix of WT KPPR1 and KPPR1ΔgltA. Bacterial burden in the blood, spleen, liver, and lung was measured after 24 hours, and log competitive index of the mutant strain compared to the WT strain was calculated for each mouse strain (n = 9-11/group, mean displayed, \**P* < 0.05, \*\*\*\**P* < 0.00005, one-sample *t* test or Student’s *t* test). (B) C57BL/6J mice or isogenic *Lcn2-/-* mice were orally inoculated with approximately 5×10^6^ CFU of a 1:1 mix of WT KPPR1 and KPPR1ΔgltA. After 48 hours, mice were euthanized and cecal bacterial load was measured by dilution plating (n = 12-17, mean displayed, \**P* < 0.05, Student’s *t* test).

## Discussion

The ability of bacteria to utilize available nutrients upon encountering a new environment, referred to as metabolic flexibility, is critical for its survival and success. In order to achieve this end, bacteria must control highly interconnected metabolic pathways that are quickly activated based substrate availability in their local environment. Central carbon metabolism connects all pathways in the cell by providing carbon skeletons for biosynthesis of macromolecular building blocks and conversely represents convergence points for the catabolism of macromolecules. Central carbon metabolism is comprised of glycolysis, gluconeogenesis, the Entner-Doudoroff pathway, the pentose phosphate pathway, and the citric acid cycle. The research presented here identifies the citric acid cycle component, citrate (Si)-synthase *(gltA)*, as a critical mediator of metabolic flexibility in Kp, and this metabolic flexibility drastically influences fitness during infection in a site-specific manner. Using multiple murine models of infection in *Lcn2+/+* and *Lcn2-/-* backgrounds, we show that *gltA* is a fitness factor during lung infection by direct and hematogenous routes, but not necessary for bacteremia. Additionally, *gltA* is necessary for gut colonization, which frequently precedes infection [12,13]. The necessity of *gltA* is determined by the nutrient composition of each respective body site, more specifically, access to amino acids that the bacteria cannot synthesize *de novo* without GltA. This is supported by the observation of differential fitness of KPPR1ΔgltA in the *Lcn2+/+* and *Lcn2-/-* lung, which have different endogenous levels of amino acids. Together, these data provide new insight into how Kp metabolic flexibility determines fitness during infection. The impact of this finding is highlighted by the fact that *gltA* is highly conserved among Kp strains. While the contributions of specific metabolic processes, such as iron acquisition [24,25,35,36,45–48], nitrogen utilization through urease activity [49], allantoin metabolism [50], and psicose metabolism [51] in Kp pathogenesis have been explored, this study is the first to reveal a role for central metabolism during Kp infection.

GltA is a type II citrate synthase, which are characteristically found in Gram-negative bacteria. The Kp GltA is closely related to the citrate synthases of other members of *Enterobacteriaceae*, such as S. *enterica* and *E. coli*, sharing 96% and 95% amino acid sequence identity, respectively. Interestingly, the Kp genome has multiple genes annotated as *gltA.* Apart from VK055_1802, VK055_2057 is annotated as *gltA* (hereafter *gltA2).* The *gltA2* gene is universally present in *Klebsiella spp.* but not in other *Enterobacteriaceae* genera. I-TASSER 3D structure prediction [52–54] indicates that GltA2 is structurally similar to GltA despite sharing only 60% and 58% amino acid sequence identity with KPPR1 GltA and *E. coli* str. K-12 substr. MG1655 GltA, respectively. Despite this structural similarity, our data demonstrate that *gltA* and *gltA2* were functionally distinct, as the presence of *gltA2* was unable to complement the loss of *gltA.* While the impact of *gltA2* on metabolic flexibility has yet to be explored, it is beyond the scope of this study; however, the universal presence of *gltA2* in *Klebsiella sp.* but not in other *Enterobacteriaceae* may provide a unique opportunity as a target for species and sub-species level detections.

Our data indicate that the loss of *gltA* abrogates the *de novo* synthesis of L-glutamate and other key carbon skeletons for biosynthesis of amino acids, which is essential to bacterial survival. Glutamate is needed for ammonia assimilation and is the fulcrum of glutamine, proline, arginine, and histidine biosynthesis [55]. In fact, glutamate and glutamine provide nitrogen for all nitrogen-containing components of the bacterial cell, and approximately 88% comes from glutamate [56]. Studies in *E. coli* have shown that glutamate is the most abundant intracellular metabolite, with an absolute intracellular concentration of 96 mM [57]. Additionally, after conversion to its D-enantiomer, glutamate serves as a component of bacterial peptidoglycan, which forms the cell wall and determines the rate of cell elongation [58]. Thus, Kp lacking *de novo* synthesis of glutamate are incapable of proliferating unless glutamate can be acquired exogenously. This is highlighted by the fact that growth of KPPR1ΔgltA in M9 is rescued by the addition of glutamate in a dose-dependent manner. A complete restoration of growth only occurs upon the addition of a sufficient amount of glutamate (Fig 3). Alternatively, exogenous glutamine, proline, arginine, and histidine can facilitate growth through the central nitrogen metabolic circuit or through production of glutamate through degradation [55]; however, supplementation of 2-oxoglutaric acid at a high concentration was not sufficient for a complete restoration of growth, suggesting a lack of transporting mechanism (Fig S2). Additionally, our data demonstrate that dipeptides containing glutamate, glutamine, and histidine can support growth of GltA-deficient Kp (Fig 2), suggesting that bacteria rely on scavenging dipeptides or polypeptides in the course their colonization and infection. Taken together, our findings suggest that stratifying *in vivo* environments as either nutritionally replete or deplete relative to the bacteria is apropos, and that a systems biology approach of studying bacterial metabolic flexibility is beneficial for understanding the lifestyle of pathogenic bacteria.

By maintaining high metabolic flexibility, pathogenic bacteria can invade multiple niches, and thus, increase their chances of evolutionary success. For example, commensal *E. coli* living in the human gut favor metabolic pathways that take advantage of the sugar-rich mucus lining [59,60], whereas uropathogenic *E. coli* (UPEC) favor metabolic pathways that take advantage of the nitrogen-rich urinary tract environment [61]. As predicted, deletions in glycolysis, pentose phosphate pathway, and the Entner-Doudoroff pathway had little effect on fitness in the urinary tract environment, whereas specific citric acid cycle components are necessary [9,62]. Similarly, *Legionella pneumophila* utilizes amino acids as its primary carbon source during lung infection in immunocompromised patients [63]. Our data suggests that metabolic flexibility plays a similar role for Kp as it does for *E. coli*, wherein the ability to utilize mucin sugars is important for survival in the gut and the ability to utilize peptides and amino acids is important for Kp survival during lung infection. As abrogation of glycolysis, pentose phosphate pathway, and Entner-Doudoroff pathway enzymes do not impact UPEC survival in the urinary tract, one would expect that abrogation of these pathways has little effect on Kp fitness in the lung. Indeed, transposon interruptions of specific glycolysis *(pfkB*, VK055_3061 [pgi], *tpiA, pyk)*, pentose phosphate pathway (gnd), and Entner-Doudoroff pathway (VK055_1337 *[tal1]*, VK055_2566 [*tal3*], *edd*) enzymes were not functionally complemented in the *Lcn2-/-* lung (Dataset S1). Correspondingly, one would predict that specific citric acid cycle components are important for Kp fitness in the lung. Interestingly, only two citric acid cycle enzymes, *gltA*, and the fumarate reductase subunits, *frdD*, were functionally complemented in the *Lcn2-/-* lung; however, the fumarate reductase subunits *(frdA-C)* did not display a similar phenotype (Dataset S1), and furthermore, these enzymes are only used during anaerobic growth. Our previous InSeq study identified the citric acid cycle components *gltA, frdA*, and *frdC*, as fitness factors during lung infection [26]. Accordingly, only one glycolytic enzyme *(pfkB)*, no pentose phosphate pathway enzymes, and no Entner-Doudoroff pathway enzymes were identified as fitness factors in this study [26]. Surprisingly, interruption of the citric acid cycle genes *acnA* and *frdB* and the Entner-Doudoroff pathway enzyme *tal1* resulted in enhanced fitness during lung infection in these studies; however, the underpinnings of this phenotype have yet to be explored. Taken together, these findings indicate that the oxidative citric acid cycle is beneficial for Kp lung infection, though under specific nutrient conditions only parts of the cycle are necessary for complete fitness. Furthermore, our findings highlight the importance of *gltA* to the metabolic flexibility of Kp during niche invasion in the human host and lay the groundwork for more comprehensive studies aimed at understanding the metabolic requirements for invasion of different body sites and the complex interactions between different metabolic pathways in the context of infection.

Phenotypic metabolic flexibility has also been used to delineate closely related species of the *K. pneumoniae* complex, which includes *K. pneumoniae, K. quasipneumoniae*, and *K. variicola*, as well as Kp pathogenic lineages. *K. quasipneumoniae* is largely considered to be an opportunistic pathogen that is frequently found as a colonizer [64], whereas *K. variicola* causes more serious infections [65]. The three members of this complex can be separated by their metabolic profile, wherein metabolism of adonitol, psicose, tricarballylic acid, and hydroxyproline phenotypically separates Kp from *K. quasipneumoniae* and *K. variicola* [66]. Moreover, the Kp strain used in this study, KPPR1, has been shown to be more metabolically flexible than the less pathogenic Kp strain MGH 78578 [67]. Finally, metabolism of D-arabinose [66] and allantoin [50] is associated with hyper-virulent Kp strains. As such, metabolic flexibility may be a critical dictator of the variation in clinical outcomes for different *Klebsiella spp.* or Kp pathogenic lineages. Our data, which support the above literature (Dataset S2, Fig S4), suggests that this is indeed the case, as reduction of metabolic flexibility through deletion of *gltA* drastically modifies fitness in different body sites (Figs 1, 6, S4). Subtle fine-tuning of metabolic capacity may confer virulence or hypervirulence to a relatively avirulent Kp strain. Indeed, *gltA* has previously been implicated to play a significant role in metabolic fine-tuning in the Lenski Experiment, wherein mutation of *gltA* permitted access to a novel nutritional niche [68]. Further exploration of the determinants of metabolic flexibility in Kp and the respective association with clinical outcomes is necessary to fully understand how specific metabolic capacity influences fitness during infection.

While this study significantly advances our understanding of the role that metabolic flexibility plays in determining fitness during infection, it is not without its limits. Firstly, this study does not address the mechanism underlying the difference in amino acid content between the *Lcn2+/+* and *Lcn2-/-* lung. Initially this result was unexpected; however, the effects of Lcn2 are not limited to antimicrobial activity. Lcn2 has the ability to act as a growth and differentiation factor [69,70], as well as the ability to modulate expression of many lung epithelial cell genes [25]. While the role of Lcn2 in these processes is not well understood, it may be the case that deletion of *Lcn2* impacts lung homeostasis, leading to altered physiology that explains the increase in amino acid levels. Although understanding the mechanism underlying this phenotype is beyond the scope of this study, the phenotype served as a useful tool to observe the effect of increased amino acid levels on Kp lung fitness. Secondly, this study exclusively uses the Kp strain KPPR1. Thirdly, we have focused solely on the citrate (Si)-synthase GltA in describing the role of metabolic flexibility during Kp infection and have not included any additional metabolic enzymes. Additional studies including different strains of the *K. pneumoniae* complex and focusing on additional metabolic pathways are necessary to fully understand the impact of metabolic flexibility during infection.

In summary, we have described a novel role the Kp citrate synthase gene, *gltA*, as a critical mediator of site-specific fitness during infection due to its influence on metabolic flexibility. Taken together, our results represent an advancement in our understanding of Kp metabolism during infection and enhance our knowledge of how these serious infections manifest, such that we are better able to combat these dangerous bacteria.

## Materials and Methods

### Ethics statement

This study was performed in strict accordance with the recommendations in the *Guide for the Care and Use of Laboratory Animals* [71]. The University of Michigan Institutional Animal Care and Use Committee approved this research (PRO00007474).

### Materials, media, and bacterial strains

All chemicals were purchased from Sigma-Aldrich (St. Louis, MO) unless otherwise indicated. *K. pneumoniae* KPPR1 [27], which is referred to as “KPPR1” throughout this study, and isogenic mutants were cultured in Luria-Bertani (LB, Becton, Dickinson and Company, Franklin Lakes, NJ) broth at 37°C with shaking, or in M9 minimal medium (M9 salts [Thermo Fisher Scientific, Waltham, MA], 0.2 M MgSO_4_, 0.01 M CaCl_2_, 20% glucose) at 37°C with shaking, or on LB agar at 30°C (Thermo Fisher Scientific). The isogenic *gltA* mutant was constructed as previously described [26,31]. Briefly, electrocompetent KPPR1 cells containing a modified pKD46 plasmid encoding a spectinomycin resistance cassette [26,31] were electroporated with a gltA-specific target site fragment containing a kanamycin resistance cassette isolated from the pKD4 plasmid [31]. Transformants were selected at 37°C on LB agar containing 25 μg/ml kanamycin, re-cultured, and confirmed by colony PCR using flanking primers (Table S1). The *gltA* complement plasmid was constructed using a Gibson Assembly Cloning Kit (New England Biolabs, Ipswich, MA). Briefly, the *gltA* sequence including its promoter was amplified from WT KPPR1 by PCR (Table S1) and ligated into the pACYC184 backbone [72] to create the pGltA plasmid. The ligation mixture was transformed into NEB 10-beta Competent *E. coli* (New England Biolabs) by heat shock. Transformants were selected at 37°C on LB agar containing 30 μg/ml chloramphenicol, re-cultured, and confirmed by colony PCR using (Table S1). Singe transformants were then grown in batch culture for plasmid extraction using the Plasmid Midi Kit (Qiagen, Germantown, MD). KPPR1ΔgltA competent cells were prepared as previously described [26], electroporated with the pGltA plasmid, and selected at 37°C on LB agar containing 30 μg/ml chloramphenicol. Following selection, transformants were re-cultured, and confirmed by colony PCR (Table S1) and by growth in M9 minimal broth.

### Transposon library construction and InSeq

Construction of the transposon library used in this study has been extensively described elsewhere [26]. Briefly, the pSAM_Ec plasmid [73] was modified by replacement of the ampicillin resistance cassette with a chloramphenicol acetyltransferase gene from the pKD3 plasmid [31] to create the pSAM_Cam plasmid [26]. The transposon library was constructed by mating KPPR1 with *E. coli* S17 λpir carrying the pSAM_Cam plasmid, followed by induction of the transposase by growth in the presence of 250 μM IPTG (Invitrogen, Carlsbad, CA) and selection on LB agar containing 25 μg/ml kanamycin and 30 μg/ml rifampicin. Insertion sequencing was performed as previously described [26] and deposited in the Sequence Read Archive (accession number to follow). KPPR1 has 5191 predicted genes [27], and previous calculations indicate that an inoculum of 1.1×10^5^ CFU should result in a 99% probability of each transposon mutant being present at least once during lung infection [26]. Following infection, total recovered transposon mutants were collected, gDNA was isolated using the DNeasy Blood and Tissue Kit (Qiagen, Germantown, MD), and genomic sequences adjacent to insertion sites were amplified by PCR. Following amplification, Illumina sequencing adapters were ligated to amplified junction DNA fragments, and then fragments were sequencing on an Illumina HiSeq2500 Instrument (Illumina, San Diego, CA). Sequencing reads were filtered, mapped and normalized as described [26].

### Murine models of infection

Six-to 12-week-old C57BL/6J mice *(Lcn2+/+*, Jackson Laboratory, Jackson, ME) or isogenic *Lcn2-/-* mice [28] were used for all murine models of infection. Kp was cultured overnight in LB, then bacteria were pelleted, resuspended, and diluted in sterile phosphate-buffered saline (PBS) to the appropriate dose. For lung infection studies, a dose of 1×10^6^ CFU was determined to provide sufficient representation of all mutants in the transposon library [26]. At the time of inoculation, mice were anesthetized with isoflurane, approximately 1×10^6^ CFU was inoculated in the pharynx, and the mouse was monitored until ambulatory. For validation of InSeq findings, WT KPPR1 and KPPR1ΔgltA were cultured overnight in LB, then bacteria were pelleted, resuspended, mixed 1:1, diluted in sterile PBS to the appropriate dose, and 1×10^6^ CFU was inoculated in the pharynx. After 24 hours, mice were euthanized by CO_2_ asphyxiation and lungs were collected, weighed, and homogenized in sterile PBS, and homogenates were dilution plated on selective media to determine bacterial load. For bacteremia studies, WT KPPR1 and KPPR1ΔgltA were cultured overnight in LB, then bacteria were pelleted, resuspended, mixed 1:1, diluted in sterile PBS to the appropriate dose, and mice were inoculated intraperitoneally with approximately 5×10^5^ CFU. After 24 hours, mice were euthanized by CO_2_ asphyxiation and blood, spleen, liver, and lungs were collected. Solid organs weighed and homogenized in sterile PBS, and whole blood and solid organ homogenates were plated on selective media. For oral inoculation studies, WT KPPR1 and KPPR1ΔgltA were cultured overnight in LB, then bacteria were pelleted, resuspended, mixed 1:1, diluted in sterile PBS to the appropriate dose, and mice were inoculated orally with approximately 5×10^6^ CFU. After 48 hours, mice were euthanized by CO_2_ asphyxiation and cecal contents were collected, weighed, and homogenized in sterile PBS, and homogenates were dilution plated on selective media to determine bacterial load. In all models, mice were monitored daily for signs of distress (hunched posture, ruffled fur, decreased mobility, and dehydration) and euthanized at predetermined timepoints, or if signs of significant distress were displayed. No blinding was performed between experimental groups.

### Preparation of recombinant human lipocalin 2 protein and Lcn2 growth assay

Human lipocalin 2 was recombinantly expressed, purified, and validated as previously described [46,74,75]. WT KPPR1 and various isogenic mutants were grown overnight in LB, then inoculated in RPMI with 10% (v/v) heat-inactivated resting human serum with or without 1.6 μM purified recombinant human Lcn2 at a concentration of 1 × 10^3^ CFU/mL. Cultures were incubated overnight at 37°C with 5% CO_2_, and bacterial density was enumerated by dilution plating.

### BioLog Phenotype MicroArray analysis

BioLog Phenotype MicroArrays (Biolog, Hayward, CA) analysis was performed in accordance with manufacturer’s instruction with some modifications. WT KPPR1 and KPPR1ΔgltA were cultured overnight in LB, then bacteria were pelleted, washed once in sterile PBS, then re-suspended in sterile PBS. Each strain was diluted in IF-0 medium to a final OD_600_ of 0.035, then diluted again to final inoculation concentrations as per manufacturer’s instruction. The final inoculum (100 μL) was plated onto plates PM1, PM2, and PM3. Sodium pyruvate (Thermo Fisher Scientific) was used as a carbon source for PM3 at a final concentration of 2 mM in accordance with previous metabolic phenotype analysis [66]. After inoculation, plates were sealed to avoid cross contamination of volatile compounds produced during Kp growth [66,76] and statically incubated overnight at 37°C. Following overnight incubation, growth was measured at OD_595_.

### Growth curves

The WT KPPR1 and KPPR1ΔgltA strains were cultured overnight in LB broth, then diluted to a uniform OD_600_ of 0.01 in the culture medium of interest the following day. Culture media included LB, M9 minimal medium with or without amino acid, carbon source, or serum supplementation, RPMI with 10% (v/v) serum (see “Preparation of recombinant human lipocalin 2 protein and Lcn2 growth assay” for details), and BALF (see below for details). Amino acids were supplemented at 10 mM unless otherwise indicated, and carbon sources were supplemented at 5 mg/mL. Cultures were incubated at 37°C and OD_600_ readings were taken every 15 min using an Eon microplate reader with Gen5 software (Version 2.0, BioTek, Winooski, VT) for up to 24 hours.

### Serum growth assay

The WT KPPR1 and KPPR1ΔgltA strains were cultured overnight in LB broth with antibiotic supplementation, if necessary. Bacteria were then washed with M9 minimal media by centrifugation, resuspended, and diluted to an OD_600_ of 0.01 in M9 minimal media supplemented with 20% (v/v) murine or discarded human serum. For some experiments, sera were heat-inactivated at 56 °C for 30 minutes. Cultures were grown at 37°C and OD_600_ readings were taken every 15 min using an Eon microplate reader with Gen5 software (Version 2.0, BioTek, Winooski, VT) for up to 24 hours.

### BALF growth assay

Six-to 12-week-old C57BL/6J mice *(Lcn2+/+*, Jackson Laboratory, Jackson, ME) or isogenic *Lcn2-/-* mice [28] were used for BALF collection. Briefly, mice were euthanized by CO_2_ asphyxiation and tracheas were exposed. A small incision was made in the trachea, and polyethylene tubing (external diameter 0.965 mm, internal diameter 0.58 mm, BD, Franklin Lakes, NJ) attached to a 23-gauge luer-stub adaptor and syringe containing 2 mL sterile PBS. Following tubing insertion, 4-0 silk suture (Ethicon, Somerville, NJ) was used secure the trachea and then the lungs were flushed with PBS. BALF was kept on ice until processing, wherein BALF was centrifuged at 21,130 × *g* for 30 min at 4°C to pellet contaminating bacteria, then supernatant was stored at −80°C. Rifampin was added to BALF to a final concentration of 30 μg/ml immediately prior to use. The WT KPPR1 and KPPR1ΔgltA strains were cultured overnight in LB broth with antibiotic supplementation, if necessary. Bacteria were washed with PBS by centrifugation, resuspended, diluted to an OD_600_ of 0.1 in sterile PBS, then mixed with BALF from independent mice at a ratio of 1:9. Cultures were incubated at 37°C and OD_600_ readings were taken every 15 min using an Eon microplate reader with Gen5 software (Version 2.0, BioTek, Winooski, VT) for up to 24 hours.

### Metabolomic analysis of BALF

For metabolomic analysis 100 μL of BALF was used. 20 μL was set aside to create a pooled sample from all study samples, and the remaining 80 μL of each BALF sample was prepared for GC-MS analysis according to manufacturer instructions using the Phenomenex EZFaast Free Amino Acids Analysis GC-MS kit (Phenomenex, Torrance, CA, USA). Briefly, BALF samples were combined with an internal standard (norvaline) and subjected to cation exchange solid phase extraction to purify amino acids from proteins, salts and other matrix components. The amino acids were then derivatized using a proprietary reagent and catalyst, the solvent is evaporated under a gentle nitrogen stream at room temperature, and finally the sample is resuspended for GC analysis. Quality control samples were prepared by pooling equal volumes of each sample and were injected at the beginning and the end of each analysis and after every 10 sample injections to provide a measurement of the system’s stability and performance as well as reproducibility of the sample preparation method. The pooled sample was treated identically to the study samples and was analyzed along with the samples for quality control purposes. Calibration standards were prepared containing all 20 proteinogenic amino acids at concentrations of 10, 25, 50 and 100 μM and were analyzed in replicate along with samples to enable absolute quantitation of amino acids.

GC-MS analysis was performed on an Agilent 69890N GC −5975 MS detector with the following parameters: a 1 μL sample was injected with a 1:15 split ratio on an ZB-AAA 10 m column (Phenomenex, Torrance, CA, USA) with a He gas flow rate of 1.1 mL/min. The GC oven initial temperature was 110°C and was increased at 30°C per minute to 320°C. The inlet temperature was 250°C and the MS-source and quad temperatures were 230° and 150°C respectively. GC-MS data were processed using MassHunter Quantitative Analysis software version B.07.00. Amino acids were quantitated as μM/L BALF using linear calibration curves generated from the standards listed above.

To generate these curves, all peak areas in samples and calibration standards were first normalized to the peak area of the internal standard, norvaline. Based on replicate analysis of biological samples, the quantitative variability for all reported amino acids using this method is <15% RSD.

### Statistical analysis

All *in vitro* experimental replicates represent biological replicates. For *in vitro* studies, except metabolomic analysis, two-tailed Student’s t-tests or ANOVA followed by Sidak’s multiple comparisons post-hoc test was used to determine significant differences between groups. For metabolomic analysis, R version 3.5 and the “gplots,” “pca3d,” and “rgl” packages were used for data visualization and the “limma” was package was used for false discovery rate correction. All animal studies except the InSeq study were replicated at least twice. Competitive indices were log transformed and a one-sample t-test was used to determine significant differences from a hypothetical value of 0 or two-tailed Student’s *t-* test was used to determine significant differences between groups. A *P* value of less than 0.05 was considered statistically significant for the above experiments, and analysis was performed using Prism 6 (GraphPad Software, La Jolla, CA). For InSeq analysis, a p-value was first calculated for each insertion using an exact Poisson test for comparing the two groups, and then the insertion-level p-values were combined using Fisher’s method [77] to obtain the statistical significance for each gene. Finally, the p-values were adjusted to control the false discovery rate (Dabney A and Storey JD. qvalue: Q-value estimation for false discovery rate control. R package version 1.43.0.). A *P* value of less than 1.3×10^−5^ was considered statistically significant.

## Acknowledgements and Funding

We would like to acknowledge Thekkelnaycke Rajendiran, Ph.D. and The Michigan Regional Comprehensive Metabolomics Resource Core at the University of Michigan School of Medicine for their assistance with the metabolomics aspects of this study. We would also like to thank Christ Alteri, Ph.D., Robert P. Dickson, M.D. and Nicole Falkowski for their insightful discussion, assistance, and critical edits. Finally, we would like to thank the University of Michigan Animal Care and Use staff for their assistance. This work was supported by funding from National Institution of Health (https://www.nih.gov/) grants AI125307 to M.A.B. and AI059722 and DK094777 to H.L.T.M. J.V. was supported by the Molecular Mechanisms of Microbial Pathogenesis training grant (NIH T32 AI007528). The work performed by the Metabolomics Core Services was supported by grant U24 DK097153 of NIH Common Funds Project (https://commonfund.nih.gov/) to the University of Michigan. The funders had no role in study design, data collection and analysis, decision to publish, or preparation of the manuscript. Y.S., J.V., P.B., V.F., H.L.T.M. and M.A.B. designed and performed the experiments. Y.S, J.V., and M.A.B. wrote and edited the manuscript. L.Z. designed and performed statistical analysis for InSeq experiments.

